# Comprehensive Cross-Population Analysis of High-Grade Serous Ovarian Cancer Supports No More Than Three Subtypes

**DOI:** 10.1101/030239

**Authors:** Gregory P. Way, James Rudd, Chen Wang, Habib Hamidi, Brooke L. Fridley, Gottfried Konecny, Ellen L. Goode, Casey S. Greene, Jennifer A. Doherty

**Affiliations:** Genomics and Computational Biology Graduate Program, University of Pennsylvania, Philadelphia, PA 19103, USA; Department of Systems Pharmacology and Translational Therapeutics, Perelman School of Medicine, University of Pennsylvania, Philadelphia, PA 19103, USA; Quantitative Biomedical Sciences, Norris Cotton Cancer Center, Geisel School of Medicine at Dartmouth, Lebanon, NH 03766, USA; Department of Epidemiology, Geisel School of Medicine at Dartmouth, Lebanon, NH 03766, USA; Department of Health Sciences Research, Mayo Clinic, Rochester, MN 55905, USA; Department of Medicine, David Geffen School of Medicine, University of California, Los Angeles, CA 90095, USA; Department of Biostatistics, University of Kansas Medical Center, Kansas City, KS 66160, USA; Department of Genetics, Geisel School of Medicine at Dartmouth, Hanover, NH 03755, USA

**Keywords:** Ovarian Cancer, Molecular Subtypes, Unsupervised Clustering, Reproducibility

## Abstract

Four gene expression subtypes of high-grade serous ovarian cancer (HGSC) have been previously described. In these studies, a fraction of samples that did not fit well into the four subtype classifications were excluded. Therefore, we sought to systematically determine the concordance of transcriptomic HGSC subtypes across populations without removing any samples. We created a bioinformatics pipeline to independently cluster the five largest mRNA expression datasets using *k*-means and non-negative matrix factorization (NMF). We summarized differential expression patterns to compare clusters across studies. While previous studies reported four subtypes, our cross-population comparison does not support four. Because these results contrast with previous reports, we attempted to reproduce analyses performed in those studies. Our results suggest that early results favoring four subtypes may have been driven by including serous borderline tumors. In summary, our analysis suggests that either two or three, but not four, gene expression subtypes are most consistent across datasets.

**CONFLICTS OF INTEREST:** The authors do not declare any conflicts of interest.

**OTHER PRESENTATIONS:** Aspects of this study were presented at the 2015 AACR Conference and the 2015 Rocky Mountain Bioinformatics Conference.

## INTRODUCTION

Invasive ovarian cancer is a heterogeneous disease typically diagnosed at a late stage, with high mortality (Kurman and Shih 2010). The most aggressive and common histologic type is high-grade serous (HGSC) (Vang *et al.* 2009), characterized by extensive copy number variation and *TP53* mutation (The Cancer Genome Atlas 2011). Given the genomic complexity of these tumors, mRNA expression can be thought of as a summary measurement of these genomic and epigenetic alterations, to the extent that the alterations influence gene expression in either the cancer or stroma.

Four gene expression subtypes with varying components of mesenchymal, proliferative, immunoreactive, and differentiated gene expression signatures have been reported in all studies of HGSC to date (Bonome *et al.* 2008; Tothill *et al.* 2008; The Cancer Genome Atlas 2011; Tan *et al.* 2013; Konecny *et al.* 2014). Two of these also observed survival differences across subtypes (Tothill *et al.* 2008; Konecny *et al.* 2014). Tothill *et al.* first identified four HGSC subtypes (as well as two other subtypes which largely included low grade serous and serous borderline tumors) in an Australian population using *k*-means clustering. Later, The Cancer Genome Atlas (TCGA) used non-negative matrix factorization (NMF) and also reported four subtypes which were labeled as: ‘mesenchymal’, ‘differentiated’, ‘proliferative’, and ‘immunoreactive’ (The Cancer Genome Atlas 2011). The TCGA group also applied NMF clustering to the Tothill data, and observed concordance with four subtypes (The Cancer Genome Atlas 2011). Konecny *et al.* applied NMF to cluster an independent set of HGSC samples and reported four subtypes, which they labeled as C1-C4 (Konecny *et al.* 2014). These subtypes were similar to those in the TCGA but a subtype classifier trained on these subtypes better differentiated survival in their own data, and in data from TCGA and Bonome *et al.* (Bonome *et al.* 2008).

Despite this extensive research in the area, work to date has several limitations. In both TCGA and Tothill *et al.*, ~8–15% of samples were excluded from analyses. A reanalysis of the TCGA data showed that over 80% of the samples could be assigned to more than one subtype (Verhaak *et al.* 2012). In more recent TCGA analyses by the Broad Institute Genome Data Analysis Center (GDAC) Firehose initiative with the largest number of HGSC cases evaluated to date (n = 569), three subtypes fit the data better than four (Broad Institute TCGA Genome Data Analysis Center 2014, 2015). This uncertainty in HGSC subtyping led us to determine if four homogeneous subtypes exist across study populations.

To comprehensively characterize subtypes, we analyze data from the five largest independent studies to date, including our own collection of samples, using a standardized bioinformatics pipeline. We apply *k*-means clustering as well as NMF to each population without removing “hard-to-classify” samples. Our goal is to rigorously assess the number of subtypes. These independent and parallel within-dataset analyses followed by cross-dataset comparison sidestep gene expression platform or dataset biases that could affect clustering if under or overcorrected. This contrasts with earlier work that pooled datasets together to identify subtypes (Tan *et al.* 2013) and ensures that subtypes identified are not induced by dataset or batch effects. We summarize each subtype’s expression patterns and comprehensively characterize correlations between subtype-specific gene expression across populations.

Our cross-population comparative analysis does not support that four HGSC subtypes exist; rather the data more strongly support an interpretation that there are either two or three subtypes. We show that the support for four subtypes observed in TCGA’s reanalysis of Tothill *et al.* (The Cancer Genome Atlas 2011) is lost when serous borderline tumors, which have very different genomic profiles and survival than HGSC (Ouellet *et al.* 2005; Bonome *et al.* 2005), are excluded before clustering. Our work also highlights the impact that a single study can have on the trajectory of subtyping research and suggests the importance of periodic histopathologic review and rigorous reanalysis of existing data for cross-study commonalities.

## MATERIALS AND METHODS

### Data inclusion

We applied inclusion criteria as described in the supplementary materials using data from the R package, curatedOvarianData (Ganzfried *et al.* 2013) and our own novel dataset (“Mayo”) (Konecny *et al.* 2014) (Supplementary Table S1). These criteria selected HGCS samples that were not duplicates from studies including at least 130 HGSC cases assayed on standard microarrays. Data from the new Mayo HGSC samples as well as other samples with mixed histologies and grades, for a total of 528 additional ovarian tumor samples, was deposited in NCBI’s Gene Expression Omnibus (GEO) (Edgar *et al.* 2002); these data can be accessed with the accession number GSE74357 (http://www.ncbi.nlm.nih.gov/geo/query/acc.cgi?acc=GSE74357). All study participants provided written informed consent, and this work was approved by the Mayo Clinic and Dartmouth College Institutional Review Boards.

After applying the unified inclusion criteria, our final analytic datasets include: TCGA (n = 499) (The Cancer Genome Atlas 2011; Broad Institute TCGA Genome Data Analysis Center 2014); Mayo (n = 379; GSE74357) (Konecny *et al.* 2014); Yoshihara (n = 256; GSE32062.GPL6480) (Yoshihara *et al.* 2012); Tothill (n = 242; GSE9891) (Tothill *et al.* 2008); and Bonome (n = 185; GSE26712) (Bonome *et al.* 2008) (Table 1). We restricted analyses to the 10,930 genes measured successfully in all five populations (Supplementary Fig. S1).

**Table 1.** Characteristics of the populations included in the five analytic data sets

|  | TCGA | Mayo | Yoshihara *et al.* | Tothill *et al.* | Bonome *et al.* |
| --- | --- | --- | --- | --- | --- |
| GEO |  | GSE74357 | GSE32062 | GSE9891 | GSE26712 |
| Platform | Affy HGU1133 | Agilent 4x44K | Agilent 4x44K | Affy HGU1133 | Affy HGU1133 |
| Population | United States | United States | Japan | Australia | United States |
| Original Sample Size | 578 | 528 | 260 | 285 | 195 |
| Analytic Sample Size<sup>b</sup> | 499 | 379 | 256 | 242 | 185 |
| Age [Mean (SD)] | 60.0 (11.6) | 62.9 (11.3) | NA | 60.3 (10.3) | 61.5 (11.9) |
| Stage |  |  |  |  |  |
| I | 10 (2%) | 7 (3%) | 0 (0%) | 11 (5%) | 0 (0%) |
| II | 17 (4%) | 11 (3%) | 0 (0%) | 8 (4%) | 0 (0%) |
| III | 351 (80%) | 275 (73%) | 202 (79%) | 178 (83%) | 146 (80%) |
| IV | 63 (14%) | 86 (23%) | 54 (21%) | 17 (8%) | 36 (20%) |
| Grade |  |  |  |  |  |
| 2 | 55 (12%) | 3 (1%) | 130 (51%) | 80 (37%) | NA |
| 3 | 386 (88%)<sup>a</sup> | 376 (99%) | 126 (49%) | 134 (63%) | NA |
| Debulking |  |  |  |  |  |
| Optimal | 325 (74%) | 287 (76%) | 101 (39%) | 132 (62%) | 89 (49%) |
| Suboptimal | 116 (26%) | 87 (23%) | 155 (61%) | 82 (38%) | 93 (51%) |

NA: Data not reported
<sup>a</sup>One sample was labeled as 'Grade 4' in TCGA470 <sup>b</sup>samples without survival data were excluded in survival analyses

### Clustering

We performed independent clustering within each dataset to avoid potential biases from different platforms or studies. As detailed in the Supplementary Methods, we identified the 1,500 genes with the highest variance from each dataset and used the union of these genes (n = 3,698) for clustering. We performed clustering within each dataset using each potential *k* from 2–8 clusters. We performed *k*-means clustering in each population using the R package “cluster” (version 2.0.1) (Maechler *et al.* 2014) with 20 initializations. We repeated these analyses using NMF in the R package “NMF” (version 0.20.5) (Brunet *et al.* 2004) with 100 different random initializations for each *k*. As done in prior studies, we calculated cophenetic correlation coefficients to select appropriate *k* for each dataset after NMF clustering with 10 consensus runs for *k* = 2 through 8.

### Identification of analogous clusters within and across studies

We performed significance analysis of microarray (SAM) (Tusher *et al.* 2001; Schwender *et al.* 2006) analysis on all clusters from each study using all 10,930 genes. This resulted in a cluster-specific moderated *t* statistic for each of the input genes (Schwender 2012). To summarize the expression patterns of all 10,930 genes for a specific cluster in a specific population, we combined gene-wise moderated *t* statistics into a vector of length 10,930. The TCGA subtype labels have become widely used in the field. To generate comparable labels across *k* and across studies, we mapped our TCGA subtype assignments back to the original TCGA labels to define reference clusters at *k* = 4 (that is, mesenchymal-like, proliferative-like, etc.). Clusters in other populations that were most strongly correlated with the TCGA clusters were assigned the same label.

### Clustering analysis of randomized data

Any clustering procedure is expected to induce strong correlational structure across clusters within a dataset even if there is no true underlying structure. However, if there is no true underlying structure, clusters across datasets are not expected to be correlated. To assess this, we used the same datasets but shuffled each gene’s expression vector to disrupt the correlative structure. We performed within and cross-study analyses of cluster identification using this set of data that were parallel to those performed using the non-randomized data.

### Assessing the reproducibility of single-population studies

We compared our sample assignments at *k* = 2 – 4 to the four subtypes reported in the Tothill, TCGA, and Konecny publications (Tothill *et al.* 2008; The Cancer Genome Atlas 2011; Konecny *et al.* 2014). Because the labels that were assigned in TCGA’s reanalysis of the Tothill data were not available, we performed NMF consensus clustering of Tothill’s data without removing LMP samples in order to generate labels for comparison.

### Data availability

We provide software to download the required data and reproduce our analyses. The software is provided under a permissive open source license (Way *et al.* 2015). Analyses were run in a Docker container, allowing the computing environment to be recreated (Boettiger 2015). Our Docker image can be pulled from here: https://hub.docker.com/r7gregway/hgsc_subtypes/. This allows interested users to freely download the software, reproduce the analyses, and then build on this work. All data used in this analysis is publicly available including data we generated (accessible under GEO accession GSE74357).

## RESULTS

### Clustering

To visually inspect the consistency and distinctness of clusters, we compared sample-by-sample correlation heatmaps. For *k* = 2 to 4 within each study, we observed high sample-by-sample correlations within clusters and relatively low sample-by-sample correlations across clusters (Supplementary Fig. S2). Clustering results using NMF were similar to *k* means results (Supplementary Fig S3).

### Correlation of cluster-specific expression patterns

Across datasets, we observed strong positive correlations of moderated *t* score vectors between analogous clusters in TCGA, Tothill, Mayo, and Yoshihara (Fig. 1; Table 2). However, clustering of the Bonome data did not correlate strongly with clusters identified in the other datasets (Table 2). We believe that we were unable to assign parallel subtypes in Bonome because of either RNA contamination or inappropriate grading assignments. However, more work is required in order to identify exactly why we were unable to classify.

**Figure 1.**
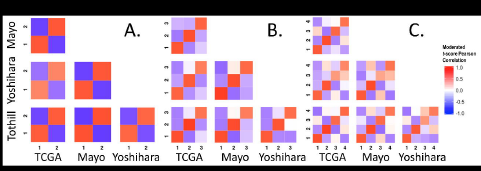
Significance analysis of microarray (SAM) moderated *t* score Pearson correlation heatmaps reveal consistency across datasets. (A) Correlations across datasets for *k* means *k* = 2. (B) Correlations across datasets for *k* means *k* = 3. (C) Correlations across datasets for *k* means *k*
= 4

**Table 2.** SAM moderated *t* score vector Pearson correlations between analogous clusters across populations^a^

|  | Cluster 1 | Cluster 2 | Cluster 3 | Cluster 4 |
| --- | --- | --- | --- | --- |
| $k = 2^a$ | 0.62 – 0.81 | 0.62 – 0.81 | NA | NA |
| $k = 3^a$ | 0.77 - 0.85 | 0.80 - 0.90 | 0.65 - 0.77 | NA |
| $k = 4^a$ | 0.77 - 0.85 | 0.83 - 0.89 | 0.51 - 0.76 | 0.61 - 0.75 |
| Bonome $k = 2^b$ | -0.08 – 0.24 | -0.08 – 0.24 | NA | NA |
| Bonome $k = 3^b$ | 0.45 – 0.46 | -0.02 - 0.12 | 0.22 - 0.42 | NA |
| Bonome $k = 4^b$ | 0.50 - 0.57 | -0.04 - 0.04 | 0.13 - 0.29 | 0.26 - 0.43 |

<sup>a</sup>Correlation ranges for TCGA, Mayo, Yoshihara, and Tothill.
<sup>b</sup>Bonome is removed from gene set analyses because of low correlating clusters

To assess our analytical approach, we performed an analysis using randomized data. This showed that within-population correlation structure was induced by clustering, but structure between populations was not (Supplementary Fig. S4). Comparing Figure 1 with S4, we observed much higher correlation across datasets (Fig. 1), which was lost after randomization (Supplementary Fig. S4). For example, for *k* = 2, the TCGA and Mayo cluster correlations for analogous clusters was high (top left panel in Fig. 1). Conversely, the same relationship in randomized data (second row, first column panel in Supplementary Fig. S4) showed correlations near zero. This indicates that the high correlations observed across datasets in Figure 1 are induced by similar underlying structure in the data.

Across studies, positive correlations between analogous clusters and negative correlations between non-analogous clusters were stronger for clusters identified when *k* = 2 and *k* = 3 than when *k* = 4 (Fig. 1), with comparable statistical precision (Supplementary Table S2). These cross-population comparisons suggested that two and three subtypes fit HGSC gene expression data more consistently than the four widely accepted subtypes.

Within each population, clusters identified by NMF were similar to those identified using *k*-means clustering (Fig. 2) suggesting that these results were independent of clustering algorithm. With NMF, both positive and negative correlations were stronger for *k* = 2 and *k* = 3 than for *k* = 4. Across *k* = 3 and *k* = 4, correlations were strongest for clusters 1 and 2. Sample cluster assignments for both *k*-means and NMF clusters are provided in Supplementary Table S3.

**Figure 2.**
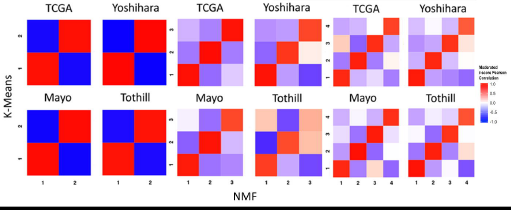
Significance analysis of microarray (SAM) moderated *t* score Pearson correlation heatmaps of clusters formed by *k* means clustering and NMF clustering reveals consistency across clustering methods. Within dataset results are shown for both methods when setting each algorithm to find 2, 3, and 4 clusters.

### Comparison with previously-identified HGSC clusters

Our clustering results for the Tothill, TCGA, and Mayo datasets were highly concordant with the clustering described in the original publications (Tothill *et al.* 2008; The Cancer Genome Atlas 2011; Konecny *et al.* 2014), as evidenced by the high degree of consistent overlap in sample assignments to the previously-defined clusters (Table 3). Our cross-study cluster 1 was mostly mapped to the “Mesenchymal” label from TCGA, “C1” from Tothill, and “C4” from Mayo. This cluster was the most stable in our analysis within all datasets, across *k* = 2, 3 and 4, and across clustering algorithms. Cross-study cluster 2, which was also observed consistently, was most similar to the “Proliferative” label from TCGA, “C5” from Tothill, and “C3” from Mayo. Cross-study cluster 3 for *k* = 3 was associated with both the “Immunoreactive” and “Differentiated” TCGA labels, “C2” and “C4” in Tothill, and “C1” and “C2” in Mayo. For analyses where *k = 4,* the third cluster was associated with “Immunoreactive”, “C2”, and “C1” while the fourth cluster was associated with “Differentiated”, “C4”, and “C2” for TCGA, Tothill, and Mayo respectively.

**Table 3.**
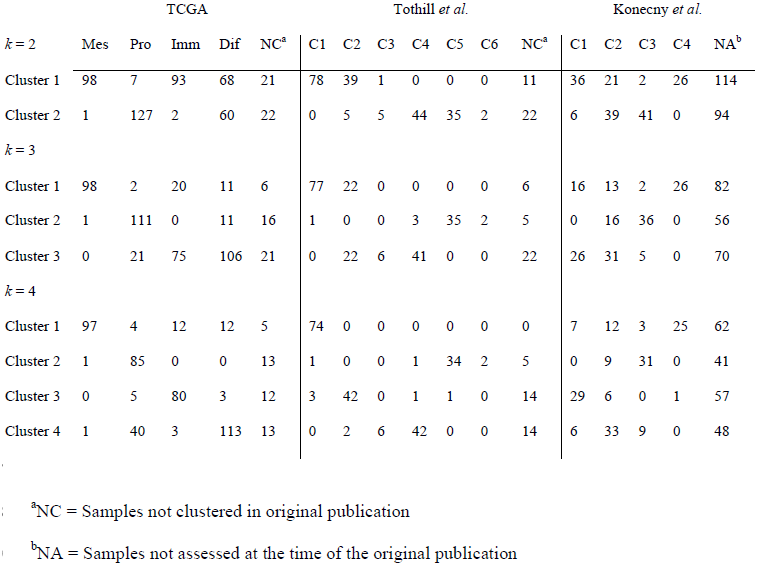
Distributions of sample membership in the clusters identified in our study by the original cluster assignments in the TCGA, Tothill, and Konecny studies. Clusters identified in our study using *k*-means clustering with *k* = 2, *k* = 3, and *k* = 4

NOTE: The corresponding labels for the generally similar HGSC gene expression subtypes observed in the TCGA, Tothill, and Konecny studies are, respectively: mesenchymal/C1/C4, proliferative/C5/C3, immunoreactive/C2/C1, and differentiated/C4/C2)

### Meta-Research into previous HGSC subtyping studies

Each of the publications that only considered high-grade samples (TCGA and Konecny *et al.*) found clustering coefficients consistent with *k* = 2, *k* = 3, and *k* = 4. Nevertheless, each publication concludes the existence of four subtypes, while our cross-population analysis suggested that two or three clusters fit HGSC data better than four clusters.

To compare with previous results, we evaluated the number of subtypes that fit the data best within each study by calculating cophenetic correlation coefficients at *k* = 2 through k=8 clusters inclusively. We observed a similar pattern in each population (Supplementary Fig S5 – S7; Fig. 3A) in which the highest cophenetic correlation was reached for two clusters and, based on the heatmaps, appeared to have the highest consensus. In every dataset, four clusters were not observed to represent the data better than two or three. The only results in previous studies that contradicted this work were from TCGA’s reanalysis of the Tothill data. According to supplemental figure S6.2 in the TCGA paper, the reanalysis included serous borderline tumors (i.e., tumors with low malignant potential) (n = 18). The inclusion of these tumors in the TCGA HGSC analyses was done even though, in the original Tothill paper, the serous borderline tumors had a unique gene expression patterns and clustered entirely in a group labeled “C3”.

**Figure 3.**
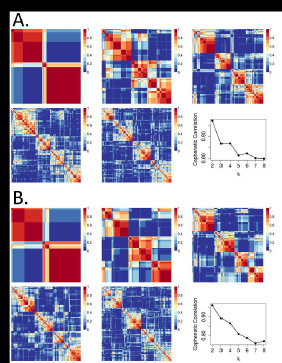
Comparing NMF consensus clustering in the Tothill dataset. Data displays consensus clustering for *k* = 2 to *k* = 6 for 10 NMF initializations alongside the cophenetic correlation results for *k* = 2 to *k* = 8. (A) Tothill dataset (n = 260) with low malignant potential (LMP) samples (n = 18) not removed prior to clustering. (B) Tothill dataset with LMP samples removed (n = 242).

To assess the extent to which serous borderline tumors inclusion drove the TCGA results, we reproduced TCGA’s reanalysis of Tothill *et al.*, including the serous borderline tumors (n = 18); we indeed observed that the cophenetic correlation is higher for *k* = 4 than *k* = 3 (Fig. 3A). However, when we appropriately removed these serous borderline tumors we observed an increase in the *k* = 3 cophenetic correlation (Fig. 3B). The results that support four subtypes were generated during clustering of HGSC and serous borderline tumors combined. Subtyping analyses of HGSC alone reveal less than four subtypes. Even after subtyping there remains a complex and nuanced portrait of the disease.

## DISCUSSION

Although prior studies have reported the existence of four molecular subtypes of HGSC ovarian cancer (The Cancer Genome Atlas 2011; Tothill *et al.* 2008; Konecny *et al.* 2014; Broad Institute TCGA Genome Data Analysis Center 2014), our analysis suggests the existence of only two or three subtypes. This conclusion is based on our observation that concordance of analogous subtypes across study populations was stronger for two or three clusters as opposed to four. Previous studies used either *k*-means or NMF clustering, and because our results contradicted prior work, we performed analyses using both of these methods. Results for each population were similar for the *k* means and NMF clustering algorithms suggesting that the clustering algorithm did not drive the observed differences.

Because cross-population comparisons suggest that two and three clusters show more consistency than four, we explored within-study heuristics (cophenetic correlation coefficients) that suggested four subtypes in previous research. The cophenetic coefficient measures how precisely a dendrogram retains sample by sample pairwise distances and can be used to compare clustering accuracy (Sokal and Rohlf 1962). While both Konecny and TCGA reported four subtypes, in both analyses *k* = 2 and *k* = 3 resulted in higher cophenetic coefficients than *k* = 4 (Konecny Figure 2A and TCGA Figure S6.1) (The Cancer Genome Atlas 2011), (Konecny *et al.* 2014). We observed the same patterns in our own reanalysis of TCGA and analysis of the expanded Mayo cohort (Supplementary Figs. S5 and S6). Yoshihara and Tothill did not report cophenetic coefficients, but our analysis of each (Supplementary Fig S7 and Fig 3A) revealed similar patterns to TCGA and Konecny.

In the previous literature, the only report to suggest that three subtypes were inappropriate was TCGA’s reanalysis of the Tothill *et al.* data (supplemental Figure S6.2 in their publication); the cophenetic coefficient dropped dramatically at *k* = 3 before recovering at *k* = 4 (The Cancer Genome Atlas 2011). Notably, TCGA’s figure legend for this supplemental result indicates that they did not remove serous borderline tumors from the Tothill data. Our analysis of Tothill *et al.* differed from TCGA’s in that we excluded serous borderline tumors and instead supports the existence of two or three subtypes. To evaluate the influence of these serous borderline tumors in the Tothill data, we repeated our analyses including serous borderline tumors, and observed a drop in the cophenetic coefficient for *k* = 3 relative to *k* = 4 (Fig. 3). This suggests that the four subtypes observed in TCGA’s analysis of the Tothill data may be due, in part, to the inclusion of serous borderline tumors.

There are several limitations to note in the HGSC data we analyzed. Given the intratumor heterogeneity that is likely to exist (Blagden 2015), our approach would be strengthened by having data on multiple areas of the tumors. Additionally, since histology and grade classification have changed over time (Silverberg 2000; Soslow 2008), it is unclear whether the populations we studied used comparable guidelines to determine histology and grade. We attempted to exclude all low grade serous and low grade endometrioid samples because they often have very different gene expression patterns and more favorable survival compared to their higher grade counterparts (Vang *et al.* 2009). While the Bonome publication specified that they included only high-grade tumors, grade is not included in the Bonome GSE26712 data set, so we were unable to determine whether the grade distribution differs from the other studies (Bonome *et al.* 2008). It is unclear why the Bonome clusters did not correspond to the clusters observed in other populations. Lack of consistency could result from a different distribution of grade or other unreported biological differences.

In summary, our study demonstrates that two clusters of HGSC, “mesenchymal-like” and “proliferative-like”, are clearly and consistently identified within and between populations. This suggests that there are two reproducible HGSC subtypes that are either etiologically distinct, or acquire phenotypically determinant alterations through their development. Our study also suggests that the previously described “immunoreactive-like” and “differentiated-like” subtypes appear more variable across populations, and tend to be collapsed into a single category when three subtypes are specified. These may represent, for example, steps along an immunoreactive continuum or could represent the basis of a third, but more variable subtype.

Our analysis also reveals the importance of critically reassessing molecular subtypes across multiple large study populations using parallel analyses and consistent inclusion criteria. New systematic approaches hold promise for the implementation of such analyses (Celik *et al.* 2016; Planey and Gevaert 2016). Our results underscore the importance of ovarian cancer histopathology, contradict the four HGSC subtype hypothesis, and suggest that there may be fewer HGSC molecular subtypes with variable immunoreactivity and stromal infiltration.

## ACKNOWLEDGEMENTS

We would like to thank Sebastian Armasu and Hsiao-Wang Chen for help with statistical analyses and data processing and Emily Kate Shea for helpful discussions.

---

**Supplementary Figure S1.**
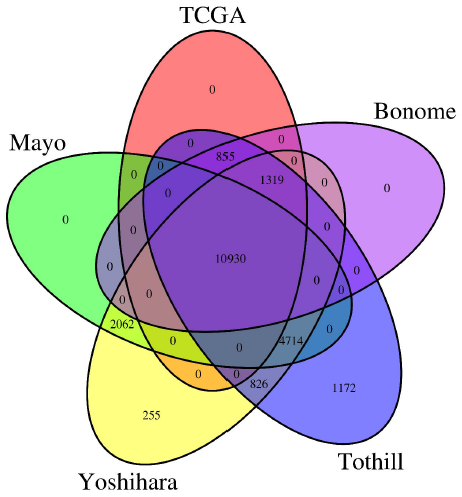
Overlapping genes assayed using either the HG-U1133 Affymetrix platform (TCGA, Tothill, Bonome) or the Agilent 4x44K platform (Mayo, Yoshihara). Differences across datasets arise from inherent array differences and/or differences in quality control preprocessing.

**Supplementary Figure S2.**
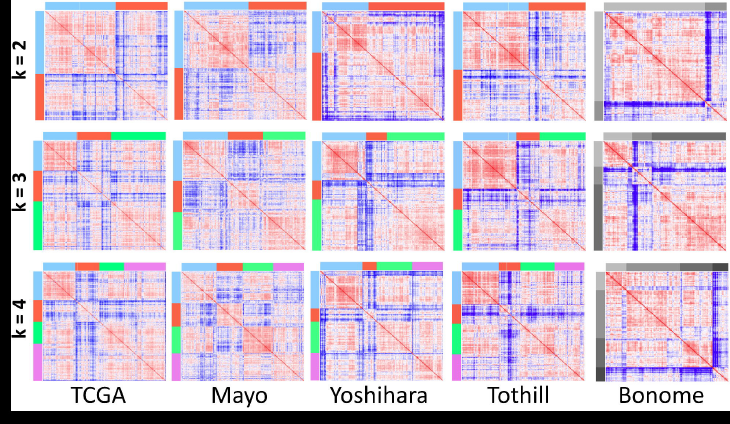
Sample by sample Pearson correlation matrices. Top panel: *k* = 2. Middle panel: *k* = 3. Bottom panel: *k* = 4. The color bars are coded as blue for cluster 1, red for cluster 2, green for cluster 3, and purple for cluster 4. In the matrices, red represents high correlation, blue low correlation, and white intermediate correlation. The scales are slightly different in each population because of different correlational structures. The clusters in the Bonome study are depicted in gray scale because in cross-population analyses to identify analogous clusters, those from Bonome did not correlate with those observed in the four other studies.

**Supplementary Figure S3.**
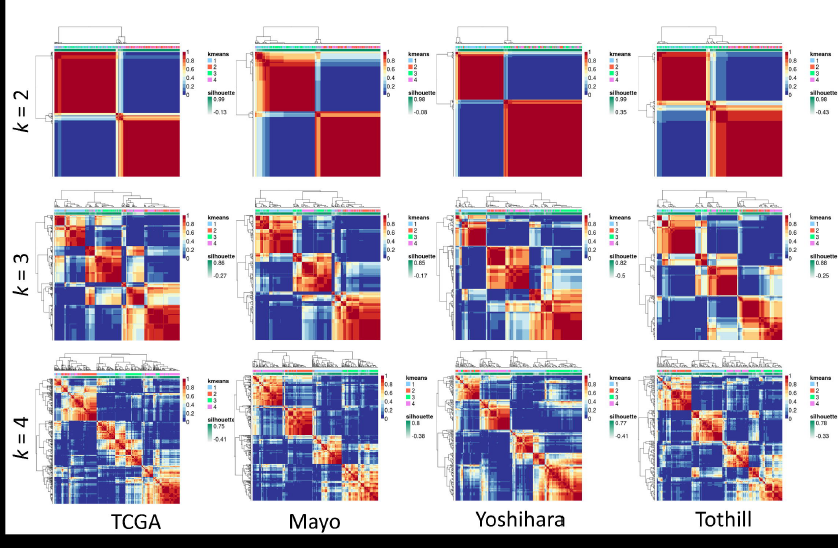
NMF consensus matrices for datasets when *k* = 2, *k* = 3, and *k* = 4. The first track represents cluster membership for *k* means clusters and the second track represents silhouette widths. Note that NMF clusters are not ordered in the same way as the *k* means clusters.

**Supplementary Figure S4.**
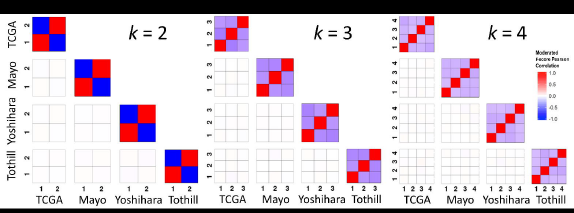
Significance analysis of microarray (SAM) moderated *t* score Pearson correlation heatmaps are not consistent across datasets for randomly shuffled gene expression values for *k* = 2, *k* = 3, or *k* = 4. The within dataset correlations are artificially induced because the clustering algorithm will find clusters even without true underlying structure. However, the across dataset clusters are not correlated in the randomized data indicating that the results we observe in Figure 1 are not artifacts of the clustering algorithm.

**Supplementary Figure S5.**
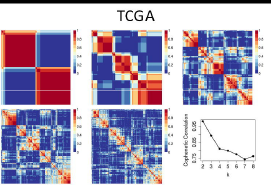
Consensus NMF clustering of the TCGA dataset (n = 499) for *k* = 2 to *k* = 6 for 10 NMF runs alongside the cophenetic correlation results for *k* = 2 to *k* = 8.

**Supplementary Figure S6.**
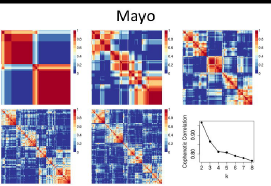
Consensus NMF clustering of the Mayo dataset (n = 379 for *k* = 2 to *k* = 6 for 10 NMF runs alongside the cophenetic correlation results for *k* = 2 to *k* = 8.

**Supplementary Figure S7.**
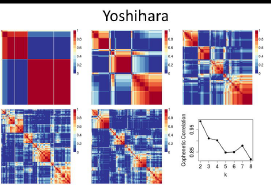
Consensus NMF clustering of the Yoshihara dataset (n = 256) for *k* = 2 to *k* = 6 for 10 NMF runs alongside the cophenetic correlation results for *k* = 2 to *k* = 8.

**Supplementary Figure S8.**
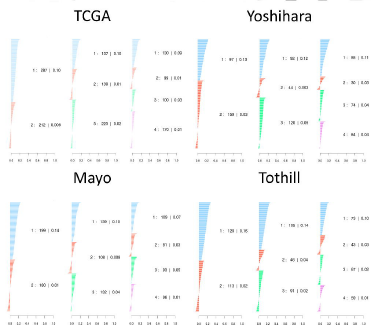
Silhouette width plots for *k* = 2, *k* = 3, and *k* = 4 for *k* means clustering results. Cluster 1 is shown in blue, cluster 2 in red, cluster 3 in green, and cluster 4 in purple.

**Supplementary Figure S9.**
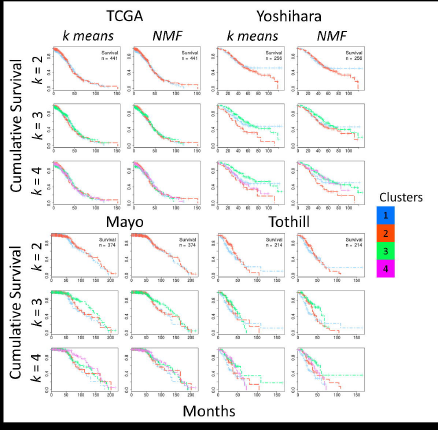
Kaplan-Meier survival curves for *k* = 2, *k* = 3, and *k* = 4 shown for clustering solutions using *k* means and NMF. Cluster 1 is shown in blue, cluster 2 in red, cluster 3 in green, and cluster 4 in purple.

